# Gain modulation and odor concentration invariance in early olfactory networks

**DOI:** 10.1101/769067

**Authors:** Emiliano Marachlian, Ramon Huerta, Fernando F. Locatelli

## Abstract

A conserved principle of the olfactory system, in most, if not all animals, is that each olfactory receptor interacts with different odorant molecules and each odorant molecule interacts with different olfactory receptors. This broad receptive field of the receptors constitutes the basis of a combinatorial code that allows animals to discriminate many more odorants than the actual number of receptor types that they express. A drawback is that high odorant concentrations recruit lower affinity receptors, which can give rise to the perception of qualitatively different odors. Here we addressed the contribution that early signal-processing in the honey bee antennal lobe does to keep odor representation stable across concentrations. We describe the contribution that GABA-A and GABA-B receptors-dependent-inhibition plays in terms of the amplitude and temporal profiles of the signals that convey odor information from the antennal lobes to the mushroom bodies. GABA reduces the amplitude of odor elicited signals and the number of glomeruli that are recruited in a concentration-dependent way. Blocking GABA-A and GABA-B receptors decreases the correlation among glomerular activity patterns elicited by different concentrations of the same odor. Based on the results we built a realistic computational model of the antennal lobe that could be further used to evaluate the signal processing properties of the AL network under conditions that cannot be achieved in physiology experiments. Interestingly, even though based on rather simplistic topology and interactions among cells solely mediated by GABA-A and GABA-B interactions, the AL model reproduced the key features of the AL stable response in relation to different concentrations.

## Introduction

The ability to recognize a stimulus or object is in itself a challenging task (Stevenson & Wilson, 2007). Different characteristics such as size, intensity, color or the background in which the object is presented can change and animals must be able to identify it regardless of those differences. This attribute is referred to as stimulus invariance and the mechanisms that make it possible vary depending on the sensory modality and on previous experience with the object (D. A. Wilson & Stevenson, 2003). In olfaction, olfactory objects are basically described at the physical world along two main dimensions; one is its chemical composition, what is normally referred to as “odor quality” and reflects the identity of the odor source, and the second one is the concentration, which is referred to as “odor intensity” and provides information in regards to the distance to the source. Importantly, at the neural level, these two dimension are not independent and it may happen that different concentrations of the same odorant are perceived as qualitatively distinct odors. In this context, odor concentration invariance, the ability of animals to recognize an odor regardless of its concentration, is studied with neurobiological interest, but also as generic theoretical problem with implications in pattern recognition technologies.

Several functional and neuroanatomical aspects of the olfactory system are repeated across animal species (Ache & Young, 2005). One of them is the fact that olfactory receptors are, in general, not specific to a unique odorant molecule but instead can interact with a wide range of different odorant molecules (Hallem & Carlson, 2006). Second, each olfactory receptor neuron (ORN) expresses only one type of olfactory receptor protein, and therefore the ORN adopts the odor tuning determined by the receptor it expresses. Third, all ORNs that express the same receptor protein send their axons to anatomically discrete subarea of the antennal lobe (AL) of insects or the olfactory bulb (OB) of vertebrates called glomeruli (Su, Menuz, & Carlson, 2009). Each odorant elicits a combinatorial pattern of active glomeruli that constitutes its specific primary representation. A drawback of this combinatorial code is that, the higher the odorant concentrations, the greater the likelihood that receptors with lower affinity are recruited into the input pattern and, therefore, different concentrations might be perceived as qualitatively distinct odors. Then, how does an animal recognize an odor along its plume, either when it approaches the odor source, or when it gets away from it? In this context, the neural mechanisms that contribute to equalize the representation of an odor across concentrations are studied insects (Froese, Szyszka, & Menzel, 2014; Luo, Axel, & Abbott, 2010; Olsen, Bhandawat, & Wilson, 2010; Olsen & Wilson, 2008; Stopfer, Jayaraman, & Laurent, 2003; Zhu, Frank, & Friedrich, 2013) and vertebrates (Arneodo et al., 2018; C. D. Wilson, Serrano, Koulakov, & Rinberg, 2017).

Here we use the honey bee antenal lobe as model system to explore the mechanisms that provide odor concentration invariance. Honey bees have rich social and individual behaviors that rely heavily on their ability to recognize innate and learned odors. In addition, several functional and anatomical aspects of the olfactory circuits in honey bees have been described, and neural activity patterns elicited by odors can be measured along the olfactory circuit thanks to well established preparations suited for electrophysiology and calcium imaging (C. G. Galizia & Kimmerle, 2004; C. Giovanni Galizia, Eisenhardt, & Giurfa, 2012; C Giovanni Galizia & Vetter, 2004). The honey bee AL consists of approximately 160 glomeruli (Robertson & Wanner, 2006). In each glomerulus, approximately 400 olfactory receptors neurons converge onto 5-6 uniglomerular projection neurons (uPNs) (Esslen & Kaissling, 1976; Kelber, Rössler, & Kleineidam, 2006; Nishino, Nishikawa, Mizunami, & Yokohari, 2009). A dense network of GABAergic local neurons interconnect the glomeruli and contributes to synchronize PNs activity and separate the representation of odors (Girardin, Kreissl, & Galizia, 2012; Meyer & Galizia, 2012; Stopfer, Bhagavan, Smith, & Laurent, 1997). Uniglomerular projection neurons connect the AL with the lateral horn and the calyces of the mushroom bodies (Figure 1A) (Abel, Rybak, & Menzel, 2001; C. Giovanni Galizia & Rössler, 2010; Kirschner et al., 2006). At the mushroom bodies each odor is encoded by a small subset of Kenyon cells that are tuned to detect specific ensembles of coactivated projection neurons (Campbell et al., 2013; Gruntman & Turner, 2013; Honegger, Campbell, & Turner, 2011; Szyszka, Ditzen, Galkin, Galizia, & Menzel, 2005). Small or partial changes in the combination of coactivated PNs produces the activation of different Kenyon cells causing the perception of a different odor (Honegger et al., 2011). In this context, it is expected that if different odor concentrations co-activate different combination of PNs, they might produce different perceptions. Therefore, it is expected that in order to keep odor perception stable across a certain concentration range, odor elicited activation patterns at the level PNs must be able to attenuate concentration differences.

**Figure 1:**
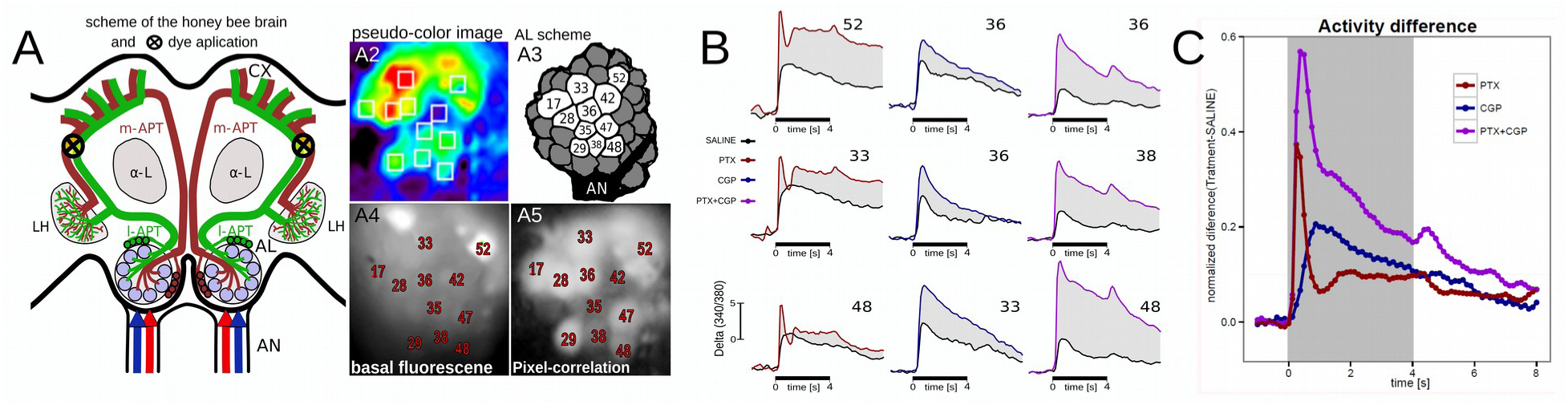
Odor-elicited calcium signals in PNs and modulation by GABA. **A**: scheme of the honey bee brain and dye injection site. Red and blue arrows indicate sensory input though the antennal nerves (AN). l-APT (green) lateral Antenno-Protocerebral-Tract, and m-APT (brown) median Antenno-Protocerebral-Tract correspond to projection neurons connecting the antennal lobes (AL) with the mushroom bodies calyces (CX) and the lateral horns (LH). α-L indicates the output lobe of the mushroom bodies that provide visual reference for the dye injection. Calcium imaging was performed on the glomeruli on the dorsal side of the ALs. **A_2_**. Representative pseudo-color image showing odor elicited activity pattern. White squares denote areas integrated as glomeruli. **A3** Antennal lobe scheme indicating the glomeruli that were identified in all individuals and therefore used for analysis. **A_4-5_**. Glomeruli were identified based on basal fluorescence after staining (380nm excitation/510nm emission), on the neighbors-pixels correlation images and by comparison with published AL atlas. **B.** Representative calcium imaging traces in identified glomeruli. Black bars at the bottom of each trace indicate stimulus duration. Odor-elicited calcium signals were measured twice in each animal. The first measurement was made under physiological saline solution (black trace) and the second measurement was made under perfusion of picrotoxin (red), CGP54626 (blue) or a cocktail of both drugs (purple). **C.** Difference between the activities measured under a given blocking condition and under physiological saline solution. Red: picrotoxin (9 bees); Blue: CGP53626 (10 bees); Purple: cocktail (11 bees). The traces represent the average of the subtraction made for every glomerulus in every animal in each condition.

The fact that several functional aspects of the input, output and local elements of the AL have been described, makes the insect antennal lobe an interesting system in which behavior, physiology and computational sciences converge to disentangle the neural mechanisms involved in sensory processing (Chan et al., 2018; Chen et al., 2015; Schmuker, Yamagata, Nawrot, & Menzel, 2011; Serrano, Nowotny, Levi, Smith, & Huerta, 2013). In this context, mathematical modeling of the cell interactions that take place in the antennal lobe allows fast evaluation of putative network configurations and also inspires solutions with technological applications in object and pattern recognition.

Here we provide a comprehensive description of gain control in odor representation in the antennal lobe. Odor elicited signals were analyzed in terms of intensity, number of recruited glomeruli and stability of the glomerular pattern that encodes the odor. This is to our knowledge, the first time that the contribution of GABA-A and GABA-B receptors dependent inhibition is systematically measured in relation to the effect of odor concentration on the glomerular patterns that encode odor identity. We found that gain control achieved by the AL network attenuates the effect of the concentration and provides a certain degree of odor concentration invariance. In addition, we generated a realistic mathematical model based on the neuroarchitecture of the honey bee AL, which replicates several features of the AL responses and allowed us to estimate the relative distribution of GABA-A and GABA-B receptors dependent inhibition across the AL network.

## RESULTS

### GABA-A and GABA-B modulate odor elicited activity in PNs

Previous studies have shown that computations in the AL optimizes the afferent input by enhancing the signal to noise contrast in odor representation, improving odor discrimination, and contributing to odor invariance across concentrations (Sachse & Galizia, 2003; Strauch, Ditzen, & Galizia, 2012). A critical element common to all these processes is the participation of local inhibitory neurons (Girardin et al., 2012; Sachse & Galizia, 2002; Stopfer et al., 1997). Here we describe the contribution that GABAergic inhibition has on shaping the output activity of the antennal lobe in honey bees. For that aim we measured odor elicited activity in the dendritic region of PNs under physiological saline solution and under perfusion with GABA-A and GABA-B receptors blockers (Farkhooi, Froese, Muller, Menzel, & Nawrot, 2013; R. I. Wilson & Laurent, 2005). PNs were backfilled with the calcium sensor dye Fura-dextran (Sachse & Galizia, 2003). Intracellular calcium increase measured by this method is mostly due to influx through voltage sensitive channels and nicotinic acetylcholine receptors (Goldberg, Grünewald, Rosenboom, & Menzel, 1999; Oertner, Brotz, & Borst, 2001) and therefore represents indirect measurement of the electrical activity (C. G. Galizia & Kimmerle, 2004).

Figure 1A shows a schematic drawing of the staining method in the honey bee brain, a representative examples of basal fluorescence of the AL after staining, neighboring-pixels correlation image used to aid glomeruli identification, pseudo-colors image showing a representative odor elicited activity pattern, and a scheme of the AL showing the glomeruli that were identified and measured in all the animals used in the present study. The traces in figure 1B correspond to representative examples of activity elicited at individual glomeruli. The black traces show odor-elicited activity measured under physiological saline solution and color traces correspond to the same glomeruli under perfusion with the GABA-A receptor blocker picrotoxin 10uM (red trace), GABA-B receptor blocker CGP54626 100uM (blue trace) or a cocktail of both blockers (purple trace). The shaded areas between the color and the black traces represents the amount of activity that in the control condition is being inhibited by GABA through GABA-A, GABA-B or both receptors together. As observed, blocking GABA receptors does not only affect the intensity of the odor elicited response but has also a drastic impact on its temporal profile. The temporal profiles of the GABA-A and GABA-B mediated inhibition were consistent across glomeruli and animals. To estimate standard temporal parameters that describe the action of GABA through the different receptors, we took every recording of every glomerulus under a given blocking condition and subtracted from it the recording obtained from the same glomerulus under control condition. The subtraction gives for every glomerulus a new trace that represents the strength of the GABAergic inhibition on a frame-by-frame resolution. Figure 1C shows the average of all those traces obtained for all glomeruli in all animals. As observed, GABA-A receptors mediate a rapid and transient inhibition of odor elicited calcium signal that peaks at 375 ms after odor onset. On the other hand, GABA-B receptors account for a predominantly tonic inhibition with a maximum inhibitory strength at 1375 ms after odor onset. When analyzing these traces it has to be considered that GABA receptor blockers do modify PNs activity not only by directly reducing inhibition to them, but also by reducing the inhibitory input into local neurons, which in turn enhances the inhibition received by PNs. It is for this reason that when PNs activity is measured in the presence of the GABA-A receptor blocker (PTX), the inhibition mediated by GABA-B receptors might be higher than its contribution in the control condition, and similarly, when activity is measured in the presence of GABA-B receptor blocker (CPG), inhibition mediated by GABA-A receptors might be also higher than its normal contribution in the control condition. In any case, it is important to take into account that GABA-A and GABA-B receptors shape activity directly and indirectly in both PNs and LNs and that GABA-A and GABA-B inhibitory components do mutually shape each other by controlling LNs activity. This may explain the rapid decay of the GABA-A inhibition until its lowest point coincident with the peak of GABA-B. In addition, the slow ascending part of the GABA-B inhibition might be explained by the early inhibition through GABA-A receptors. The contributions of GABA-A and GABA-B receptors are consistent with previous reports in *Drosophila* (Root et al., 2008; R. I. Wilson & Laurent, 2005) and cockroaches (Warren & Kloppenburg, 2014). As observed, the interplay among local neurons and the different strength with which GABA-A and GABA-B may control activity on different cell types, makes the response of this circuit a highly dynamic process. Due to this, it is very hard to predict the way the whole network contributes to equalize signals across stimulus intensities. With that question in mind, we used the data obtained so far in relation to GABA-A and GABA-B dependent inhibition to feed a realistic mathematical model of the AL that was used to test the putative distributions of inhibitory strengths that confer to the network a behavior similar to the one measured in the experiments.

### GABAergic inhibition in the AL model. Effect and validation

In order to study the effect and distribution of the GABAergic inhibition in the AL network, we built a realistic AL model that allow us to evaluate the strength and nature of the cell-to-cell inhibitory interactions that could provide encoding and signal processing properties similar to those observed in the antennal lobe. Since we know the general architecture of the honey bee antennal lobe (I. Sinakevitch, Bjorklund, Newbern, Gerkin, & Smith, 2018; I. T. Sinakevitch et al., 2013) but not the absolute connectivity of each individual neuron, neither the degree of heterogeneity across different glomeruli (Girardin et al., 2012), we opted to establish standard connectivity rules between neurons and glomeruli that vary within a minor range of possibilities that were randomly assigned. The network was composed by 20 glomeruli. Each glomerulus had 3 uniglomerular projections neurons (PNs) and 5 inhibitory local neurons (iLNs) (figure 2A). PNs acted only as post-synaptic elements that did not excite iLNs or other PNs. On the other hand, iLNs received excitatory input in only one glomerulus but could inhibit each of the 20 glomeruli with a probability of 25%. When a given iLN inhibited one glomerulus, it inhibited all neurons in it (3 PNs and 5 iLNs), except when it was in the glomerulus in which this iLN received excitatory input (home-glomerulus) in which case it inhibited the other 4 iLNs but not the 3 PNs or itself. Odor stimulation was simulated by injecting a 4 seconds pulse of depolarizing current into the 3 PNs and the 5 iLNs that formed the glomerulus. The amount of current and the number of glomeruli that were recruited could be varied to simulate different odor intensities. All neurons that were recruited by the odor received the same amount of depolarizing current. Membrane potential and ionic currents were calculated for every neuron following the Hodgkin and Huxley model (Hodgkin & Huxley, 1952) adjusted by the parameters in table 1. The temporal dynamics of the GABA-A and GABA-B dependent inhibitions obtained from the experiments in the first section were fed into the model.

**Figure 2:**
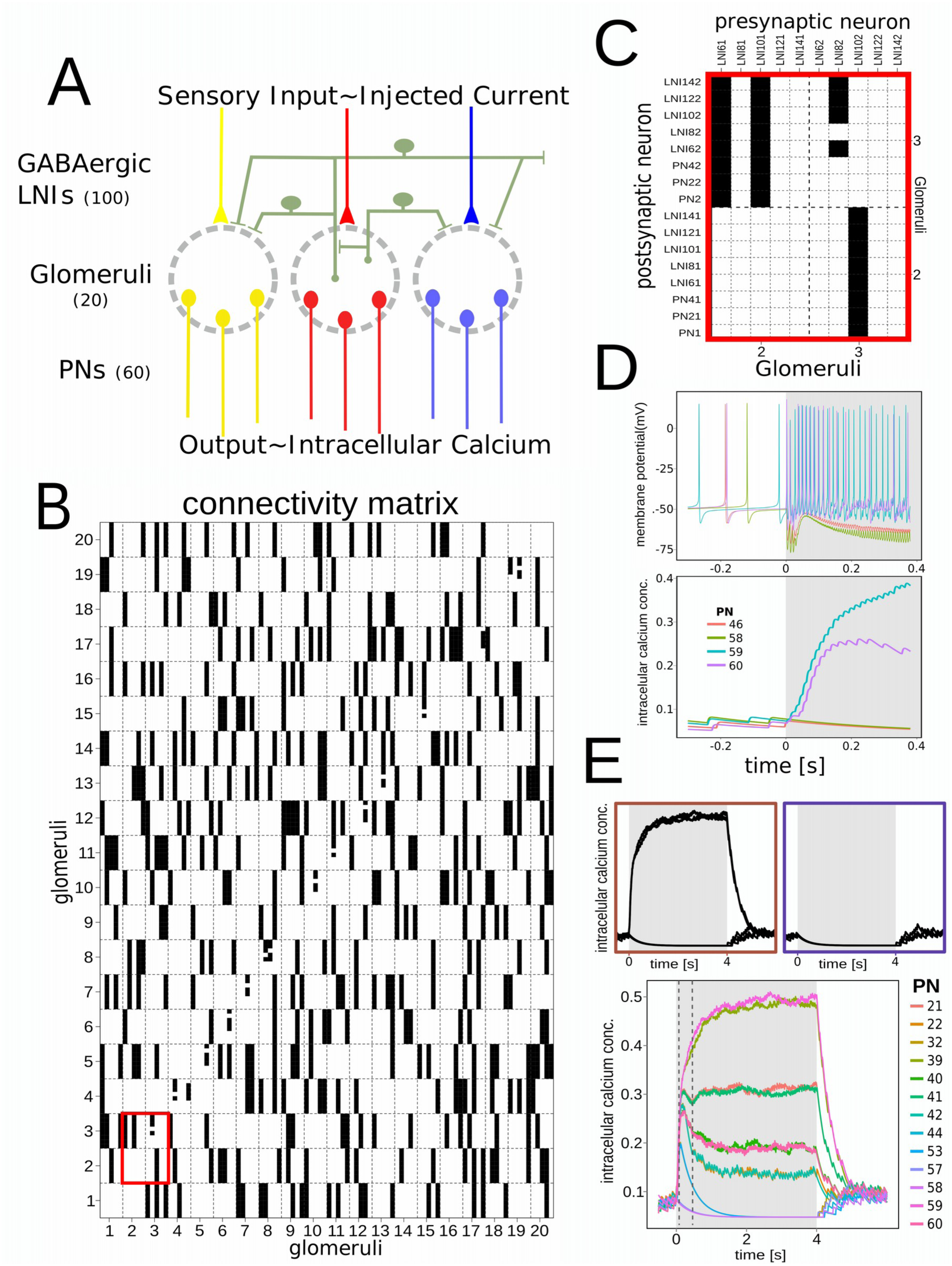
Design and connectivity of the Antennal Lobe model. A: Elements and scheme of the antennal lobe model. Yellow, red and blue components indicate input and output in three glomeruli of the 20 that comprise the AL model. Green elements represent local inhibitory neurons that interconnect glomeruli. Every local neuron receives excitatory input in one glomerulus but can inhibit every glomerulus with a probability of 25%. **B.** Random generated connectivity matrix. Every sub-column in the matrix represents one of the 100 local neurons. Local neurons are clustered in groups of 5 that receive the same depolarizing current that simulates sensory input into one glomerulus. The 20 rows represent the 20 glomeruli, each formed by 3 PNs and 5 LNs. **C**. Detail of the connectivity between 2 glomeruli. The example shows the connectivity between glomeruli 2 and 3. LNs 61 and 101 receive excitation in glomerulus 2 and inhibit all neurons (LNs and PNs) in the glomerulus 3. LNs 81, 121 and 141 receive excitatory input in glomerulus 2 but do not inhibit glomerulus 2 o 3. LN 82 receives excitatory input in glomerulus 3 and inhibits all other LNs in glomerulus 3 but not itself or the local PNs. LN102 receives input in glomerulus 3 and inhibit all neurons in glomerulus 2. LNs 62, 122 and 142 receive excitatory input in glomerulus 3 but do not inhibit glomerulus 2 or 3. **D**. Representative PNs output. Shaded area represents the stimulus input. Only four neurons are shown. Upper panel: Spontaneous firings before odor onset. Upon odor onset, two PNs increase their firing rate and two neurons are subject to inhibition. Lower panel: Intracellular calcium concentrations in the same four neurons. **E.** Three representative output patterns obtained with different strength of the inhibitory connections. Low inhibition strength: no modulation of the input is observed. High inhibition strength: the AL is silenced. With intermediate inhibitory strength, the output pattern is composed by heterogeneous responses across the PNs populations even though they have received the same excitatory input.

An example of the connectivity among the 160 neurons in the network is shown in figure 2B. Column and row numbers indicate glomeruli. The intersection among glomeruli 2 and 3 is enlarged for visualization of the connectivity between the neurons in these two glomeruli (figure 2C). A black square in the intersection between two neurons indicates that they are connected. The neurons in the columns represent the pre-synaptic partners and the neurons in the rows are the post-synaptic ones. The columns contain only iLN because the PN did not act as pre-synaptic partners of the network. When two neurons were connected, it was considered that both, GABA-A and GABA-B receptors participated of the inhibition. The model was used to test the distribution and relative strength of GABA-A and GABA-B receptors dependent inhibitions that best replicates the behavior of the real AL.

The readout of the model was based on spike frequency and intracellular calcium concentration of the PNs (figure 2D). An initial inspection of the network’s behavior showed that drastically different outputs could be generated by varying the strength of the inhibitory connections. As it might be expected, when inhibition was high, no activity was elicited at the PNs level, and when inhibition was low, the PNs output replicated the input (figure 2E). However, more importantly, we found a wide range of intermediate inhibition strengths, in which PNs reacted with a repertoire of responses, in spite of having been injected with the same amount of depolarizing current (Figure 2E, lower panel). Such heterogeneous output pattern resembles the diversity of responses that are normally elicited when stimulating olfactory circuits and can be explained in our model by the heterogeneous innervations of the local neurons implemented by the 25% random factor. The observation that the network could generate such realistic output encouraged us to further refine the strengths of GABA-A and GABA-B dependent currents that ensure a robust and rich network behavior that could mimic more reliably the network behavior observed in a real AL. The readout of the model was adapted to make it comparable to the recordings obtained in the calcium imaging experiments (Figure 1). The intracellular calcium concentrations determined for each of the three PNs in one glomerulus, were averaged to obtain a single calcium signal that represented the glomerulus response as a whole. Next, each glomerular calcium trace was binned to 8 frames per second to allow direct frame by frame comparison with the calcium imaging recordings. This procedure was made for every PN and glomerulus in the model, thus, at the end we obtained 20 calcium traces that together represented the spatio-temporal code elicited by the stimulation.

Next, we designed a procedure to evaluate to which extent the behavior of the AL model reproduces the behavior of the honey bee AL. For that aim, we took all recordings from animals in which we identified the same set of glomeruli and for each identified glomerulus we averaged across animals the response to a 4 seconds stimulation with 2-octanone (dilution 2x10-2 in liquid phase and further1/10 dilution of the headspace). The average response pattern was taken as a standard odor elicited calcium pattern. Afterwards, we calculated the Pearsońs correlation coefficient between each of the standard calcium traces and each of the 20 traces computed from the model. The highest correlation coefficient obtained was taken as the best match between a measured and a model calcium trace. This correlation value was saved and the respective pair of traces was removed to calculate the next best match between the remaining model and measured calcium traces. This procedure was repeated until having paired each of the recorded glomeruli to one of the model traces. Finally, all correlation values were averaged and this average was taken as a similarity index between the behavior of the actual AL and the model AL. This index quantifies the extent to which the response pattern generated by the AL model replicates a response pattern elicited by the real AL in terms of intensities and temporal structure of the individual traces. We used this procedure to systematically test the model while screening conditions of cell-to-cell inhibitory interactions.

Previous experiments done in fruitflies have shown that the relative contribution of GABA-A and GABA-B receptors are different depending on whether the target neuron is a LN or a PN (R. I. Wilson & Laurent, 2005). Therefore, in our AL model the strength of GABA-A and GABA-B dependent currents were considered the same across synaptic contacts of the same type but could be different between synaptic contacts of different type (LN-to-PN and LN-to-LN). We tested different strengths of: i) GABA-B dependent currents into PNs, ii) GABA-B dependent currents into iLNs, iii) GABA-A dependent currents into PNs, and iv) GABA-A dependent currents into iLNs. To screen along these multiple dimensions we first set the GABA-A dependent currents to zero and compared the output of the model with the imaging experiments performed using PTX. The color-coded matrix in figure 3A summarizes the whole search performed to find out GABA-B conditions that best reproduced the traces obtained in the PTX experiments. The *x* axis indicates the strength of the GABA-B dependent current into PNs and the *y* axis indicates the strength of the GABA-B dependent currents into iLNs. Each square corresponds to a new run of the model. The color-scale indicates the similarity index calculated between the model and the standard activity pattern. For high values of iLN-to-iLN GABA-B dependent currents, the inhibition turned off the inhibitory neurons, the input-output transformation was lost, and the PNs activity replicated the input (Figure 2E). On the other hand, when the GABA-B inhibition to PNs is high the inhibitory neurons completely turn off the PNs and the AL output is shut off (Figure 2E). As expected from a highly interconnected network, the way in which a certain level of inhibition from iLN to PN affects the output of the network was in turn affected by the strength of the inhibitions from iLN to iLN. By simultaneously screening along different inhibitory strengths of these two types of connections we found a region in which the output of the AL model resembles the response profile obtained in the calcium imaging experiments performed with PTX. Figure 3B upper panel, shows the traces generated by the simulation that provided the highest similarity index with the output pattern measured under picrotoxin (Figure 3B lower panel). As observed, the output pattern combines the features observed in the experiments, *i.e*. multiple activation levels across glomeruli, an early activation peak, an inhibition valley and the plateau.

**Figure 3:**
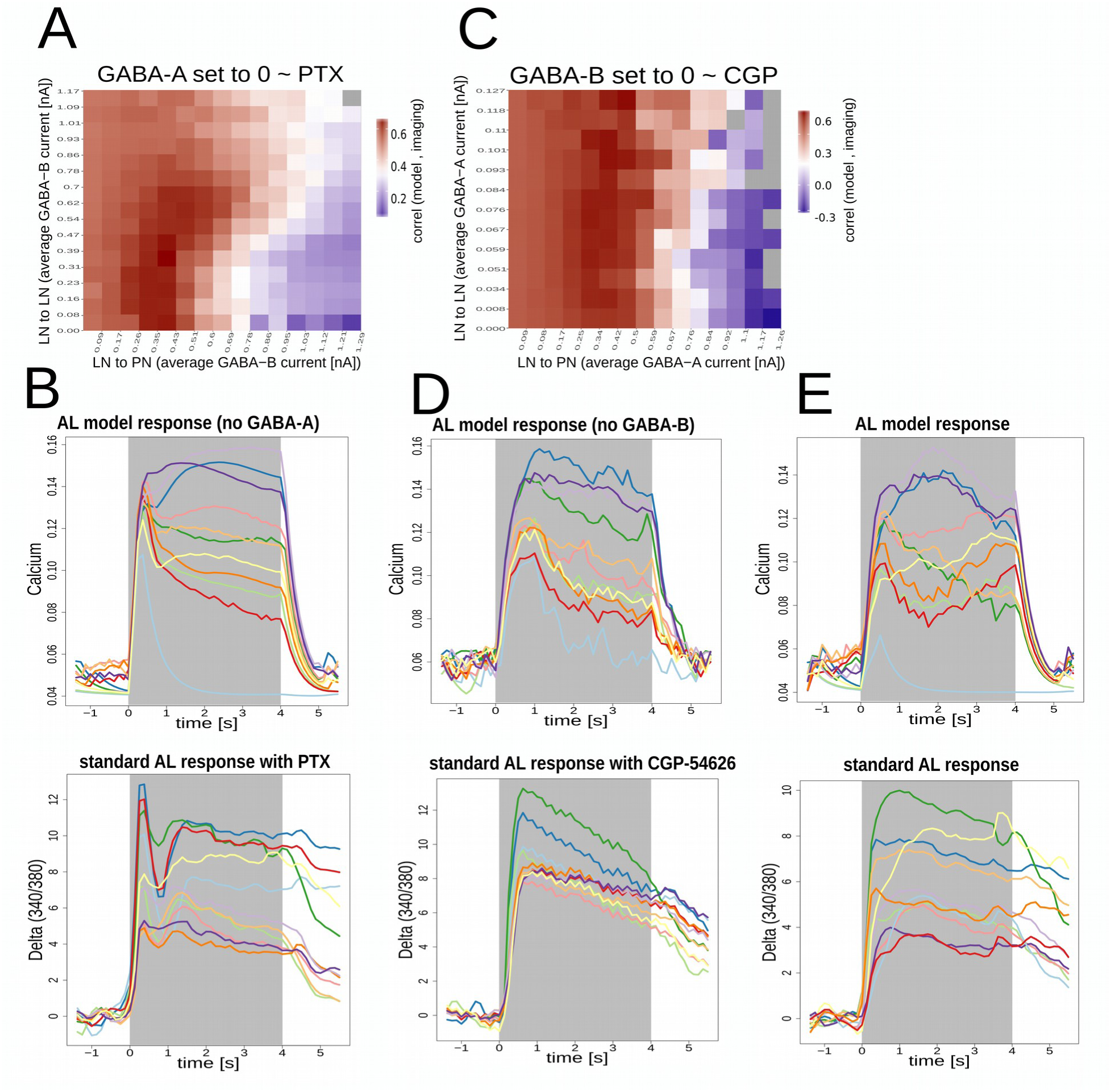
GABA-A and GABAB dependent currents. A. Search of GABA-B dependent conductances. Color matrix: similarity index between the standard AL response obtained with PTX and the model AL responses determined without GABA-A dependent currents. The strength of GABA-B dependent conductance was varied for LN-to-LN synapses and for LN-to-PN synapses to find conditions that best mimics the AL output when it was perfused with picrotoxin. Each square belongs to a new simulation. Values in the axes indicate the average inhibitory current received by each neuron (nA). **B**. Upper panel: model AL response that provided the highest similarity index with the standard AL response obtained in the presence of PTX (lower panel). **C.** Search of GABA-A dependent conductances. Color matrix: Similarity index between the standard AL response obtained with CGP56464 and the AL model responses determined without GABA-B dependent currents. The strength of GABA-A dependent conductances was varied for LN-to-LN synapses and for LN-to-PN synapses to find conditions that best mimics the Al output. **D**. Upper panel: AL model response that provided the highest similarity index with the standard AL response obtained in the presence of CGP54626 (lower panel). **E.** Upper panel: model AL response patterns obtained combining the optimal conditions determined for GABA-A dependent inhibitions into PNs and into LNs, and the optimal GABA-B dependent inhibition into PNs and LNs. Lower panel: Standard odor elicited activity pattern used as template for comparison.

Next, we proceeded in a similar way to find out the strength of the GABA-A dependent currents. We set the GABA-B dependent currents to zero and compared it with the calcium imaging recordings obtained under perfusion with the GABA-B blocker CGP54626. Varying the strength of the GABA-B dependent inhibition we found a region of parameters in which the AL output mimicked the one described in the imaging experiments. As shown in figure 3C and D, the traces generated by the model show diverse amplitudes and a temporal profile similar to the ones obtained in the measurements with the GABA-B blocker. For high values of iLN-to-PN the output did not show any activity and for high values of iLN-to-iLN currents, all PNs fired synchronously independently of the input. This last result is consistent with previous demonstrations that LNs and GABA through GABA-A receptors synchronizes PNs activity (Stopfer et al., 1997).

Finally, we enabled GABA-A and GABA-B dependent currents to simulate the fully working AL. No screening along different conductances was made this time. Instead, the model was evaluated directly using the GABA-A and GABA-B dependent conductances obtained as optimal. Considering that GABA-A and GABA-B, both inhibit PNs activity, it could have been predicted that enabling both mechanisms would intensify the inhibition of PNs. However, the output of the model reproduced the different response levels and also the complex temporal structure of the calcium traces measured under physiological saline solutions (figure 3E) replicating many response features of the honey bee AL. This result validates the values obtained in regards to the strengths with which GABA-A and GABA-B dependent inhibitions modulate LNs and PNs. It is important to emphasize the advantages provided by the mathematical modeling of the AL network which makes visible and quantifiable the complex dynamics that emerge from a highly interconnected network. These results prompted us to extend the experiments and the simulations to evaluate in which extent the GABAergic network does contribute to encode odors along different concentrations.

### GABA and gain modulation in the AL

It has been shown across species that the combinatorial pattern of glomeruli encodes odor identity and that the response intensity and the number of active glomeruli increase concomitantly with odor concentration (Julie Carcaud, Giurfa, & Sandoz, 2018; Cinelli, Hamilton, & Kauer, 1995; Falasconi, Gutierrez-Galvez, Leon, Johnson, & Marco, 2012; Friedrich & Korsching, 1997; Meister & Bonhoeffer, 2001; Ng et al., 2002; Rubin & Katz, 1999; Sachse & Galizia, 2003; Wachowiak & Cohen, 2003; Wang, Wong, Flores, Vosshall, & Axel, 2003). Despite the concentration-dependent changes in the input pattern, it is known that, unless they are trained not to, animals can generalize between different concentrations of the same odorant (Engen & Pfaffmann, 1959; Homma, Cohen, Kosmidis, & Youngentob, 2009; Masek & Heisenberg, 2008; Pelz, Gerber, & Menzel, 1997; Slotnick & Ptak, 1977; Wright & Smith, 2004). It has been suggested that this ability relies at least in part on the capability of the AL (or the olfactory bulb) to equalize odor induced signals, approximating them to a concentration invariant representation at the first olfactory interneurons. In this section we evaluated the extent in which local GABAergic inhibition contributes to the gain function at the honey bee AL output. We measured calcium signals elicited by increasing odor concentration in the dendritic region of PNs and evaluated how the different GABA blockers affect the concentration-to-signal function. 2-octanone was used as the olfactory stimulus for these experiments, because it is not consider to be a pheromone or pheromone constituent and because at id concentration it activates approximately 50% of the glomeruli at the dorsal surface of the AL. Figure 4A shows an example of the glomerular patterns and the corresponding calcium traces in identified glomeruli. The response intensity and the number of recruited glomeruli increase as odor concentration increases. In order to analyze the general intensity of the response pattern, we averaged the activity from all glomeruli and focused on the two time points in which we have previously identified the peaks of GABA-A and GABA-B dependent inhibitions, i.e. 375 ms and 1375 ms after odor onset (figure 1C). All the black traces in figure 4B show the mean and SEM of the intensity of the signals elicited upon increasing odor concentrations under perfusion with physiological saline solution. The concentration-response curve can be described by a logistic function that slows down as the odorant reaches the higher concentrations. Notice that the black traces may differ between panels, because each one corresponds to an independent set of animals. Because of this, all effects caused by the GABA blockers along the concentration series were analyzed based on within-animal comparisons of measurements performed first under control saline solution and second under perfusion with the GABA blocker. Figure 4B, left panel, shows a group of bees in which the first and second concentration series were both measured under perfusion with physiological saline solution. This group served as control for changes in calcium signals that might be caused by time or by repeated stimulation. As observed, no major difference was detected between the first and second series, neither at 375 nor at 1375 ms after odor onset (solid and dashed black lines). In three other groups of bees, the first concentrations series was measured under perfusion with physiological saline solution and the second series was made under perfusion with the GABA blockers PTX (N=11 bees), CGP54626 (N=10 bees) or a cocktail of both: PTX + CGP, (N= 9 bees). Perfusion with the GABA-A blocker PTX increased calcium signals in the time window around 375 ms but not 1375 ms after odor onset. Interestingly, the effect of blocking GABA-A was not uniform across the whole concentrations range. At low concentrations (even at zero), PTX increased signals in a concentration independent way. The gap between the black and red traces is constant from the first to the fourth concentration step. This effect likely corresponds to increased spontaneous activity after removal of a certain level of tonic GABA-A dependent inhibition. From the fifth to the eighth concentration steps, black and red traces show different slopes. This concentration dependent effect suggests a gradually growing contribution of GABA-A controlling odor elicited activity. Finally, at the three highest concentrations, the increment of the signals slows down and the difference between black and red traces remains constant, probably, as consequence of no further increase of activity at the input level. Importantly, the fact that the odor elicited calcium signals increase with PTX beyond the maximal activity reached with physiological saline solution indicates that the plateau reached by the three highest concentrations is not a ceiling artifact of the imaging method. In summary, GABA-A modulates signals across concentrations by reducing the steep function that relates odor concentration and pattern intensity.

**Figure 4.**
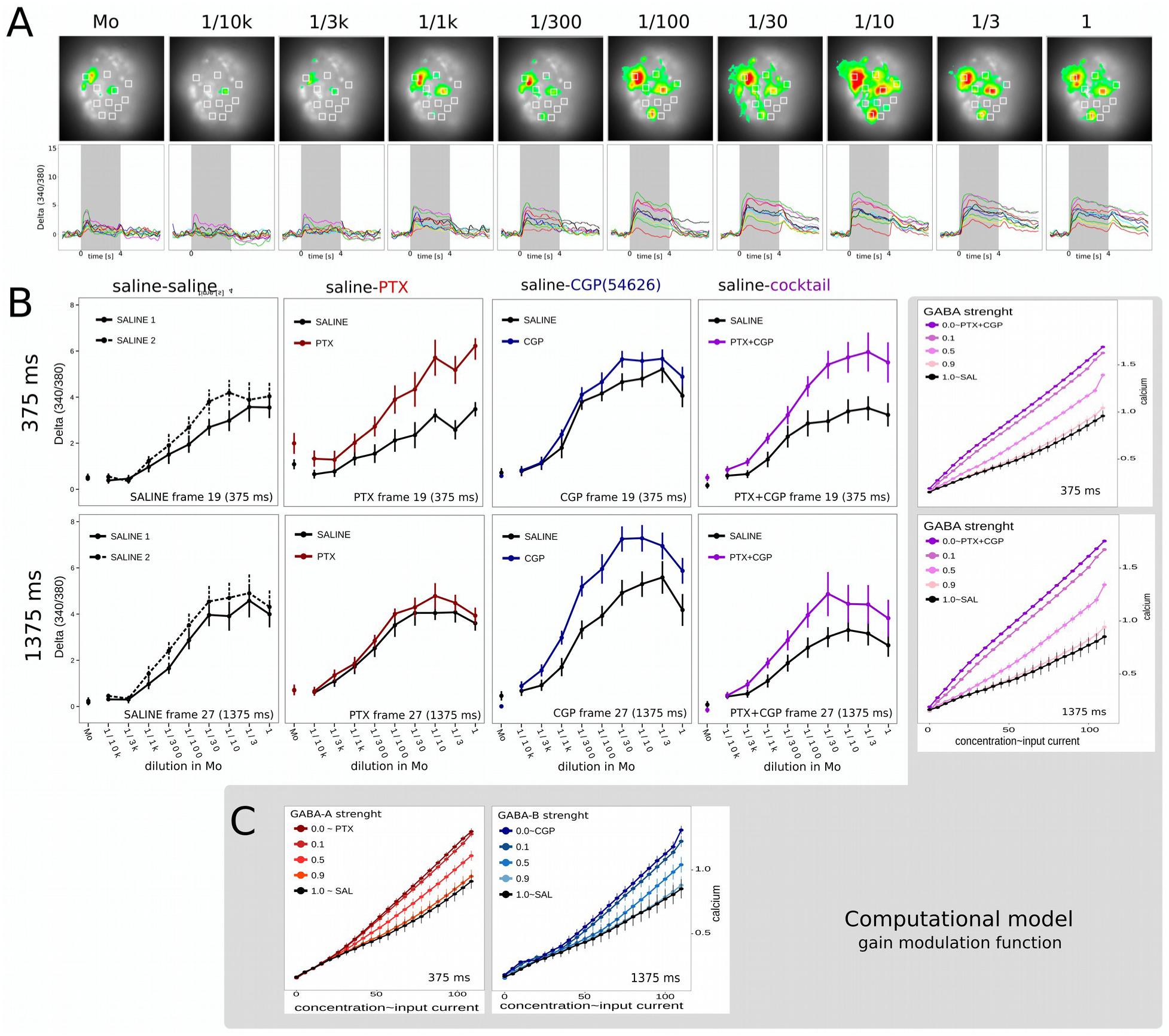
Concentration-signal function. A. Glomerular activity patterns elicited by increasing concentrations of 2-octanone. Upper panel: pseudo-color images expressing Delta(340/380) one second after odor onset. Lower panel: Temporal detail of the activity elicited in 11 glomeruli identified for the present analysis. Grey bar: stimulus duration 4 seconds. Odor concentrations: head space of 2-octanone pure, or diluted in mineral oil: pure, 1/3, 1/10, 1/30, 1/100, 1/300, 1/1000, 1/3000 and 1/10000. The head space was further diluted 1/10 in the carrier air stream before reaching the antenna. B. Four groups of bees subject to calcium imaging measurements along two concentration series each. The first group underwent two series under physiological saline solution (saline-saline) N = 8. Second group; first series saline, second series under picrotoxin 10 uM, N = 11; third group: first series saline, second series CPG54626 100 uM N = 10; fourth group: first series saline, second series cocktail picrotoxin + CGP54626 (10 uM + 100 uM), N = 9. Values represent the average Delta (340/380) across the same 11 glomeruli identified in all bees. Upper panel: activity measured 375ms after odor onset. Lower panel: activity measured 1375 ms after odor onset. C. Antennal lobe model output for increasing input currents in the AL model. Values indicate mean and SEM of the intracellular calcium in active glomeruli. In one simulation all recruited glomeruli received the same amount of depolarizing current. The amount of current changed between simulations from 0 to 100 nA. In all graphs, black series correspond to the AL output with fully working GABA-A and GABA-B dependent conductances. Red graded series correspond to gradual reduction of the strength of the GABA-A dependent currents, expressed as proportion of the highest strength (activity 375 ms after stimulus onset). Blue graded series correspond to gradual reduction of the strength of the GABA-B dependent currents (shown 1375 ms after stimulus onset). Purple graded series correspond to gradual reduction of both, GABA-A and GABA-B dependent conductances (shown both, 375 and 1375 ms after odor onset).

In another group of bees, the AL was perfused with the GABA-B blocker CGP54626 during the second concentration series (blue lines). Blocking GABA-B increased odor induced signals in the 1375 ms time window. The consequence of blocking GABA-B was clearer at the low concentrations range. Black and blue lines show different slopes from the first to the fifth concentration steps, while at higher odor concentrations, the gap between both curves remains constant. These results suggest that GABA-B may contribute to equalize signals mainly at low odor concentrations. In addition, CGP did not produce the concentration independent boost of the activity at the low concentration range that was observed with PTX. Furthermore, at zero concentration the activity with CGP was slightly lower than the activity elicited under physiological saline solution.

Finally, in the last group of bees, the first series of odor concentrations was measured under physiological saline solution (black line) and the second series was measured with a cocktail of PTX and CGP54626 (purple trace). The modulation of the gain function is observed at both time windows and along the whole concentrations range with exception of the upper region in which treatment and control responses slow down, likely because there is no further increase of sensory input beyond a certain odorant concentration.

### Gain modulation in the AL model

Next, we asked whether the signal modulation of the antennal lobe in regards to the gain modulation function can emerge from the architecture of the network and the balance of GABA-A and GABA-B inhibition strengths calculated based on the AL model.

We took the AL model as it was adjusted in the previous sections and tested it along a range of different input intensities. The input strength was controlled by the amount of injected current and fixing the percentage of recruited glomeruli to 55% (further analyzed in the last section).

First, we simulated in the model the condition in which we added PTX (figure 4C left). For that aim, the conductance associated to GABA-B dependent inhibition was fixed to the optimal value determined above and the conductances associated to GABA-A receptors were progressively reduced in successive simulation trials starting from its optimal value (black trace) to eventually reach zero conductance (orange-red-brown sequence). The general intensity of the response pattern was calculated by the average of the calcium response in all recruited glomeruli during the 375ms window. The first observation (figure 4C) is that in comparison to the figures obtained by calcium imaging, the response/input function did not saturate. This can be explained by the fact that in contrast to the real AL in which olfactory receptor neurons activity saturates at approximately 250 spikes/seconds (Hallem & Carlson, 2006), our model does not include constraints that saturate the input. An alternative explanation could be that there are other sources of inhibition which are not GABAergic and take over at higher concentrations. Leaving aside this difference, it is observed that reducing the GABA-A dependent current resembles the effect produced by PTX. The modulation in the gain function was not evident at low stimulus intensities but emerges progressively as the stimulus intensity increases. In contrast to the calcium imaging experiments, removing GABA-A did not produce the concentration-independent boost of activity at the first four concentrations steps, thus supporting the interpretation that this increase in activity may reflect spontaneous activity of sensory neurons after removal of tonic inhibition.

Next, we simulated AL conditions to test the contribution of GABA-B dependent inhibition to the gain modulation function. The conductance associated to GABA-A was set to the optimal value previously determined and the conductance associated to GABA-B was gradually reduced along successive simulation trials starting from its optimal value (black trace) to eventually reach zero conductance (light blue to dark blue sequence, figure 4C). This time we focused the analysis to the 1375ms time window. Figure 4C shows that, as the strength of GABA-B inhibition is reduced, the slope of the signal/stimulus function increases. In comparison to GABA-A, blocking GABA-B produced slight changes that, although small, were evident from very low stimulus intensities.

Finally, we tested the behavior of the AL model in relation to stimulus intensity by simultaneously reducing both, GABA A and GABA B dependent conductances (Figure 4C, right panel: purple traces). In this case, the contribution of GABA becomes evident for the whole range of stimulus intensities. When GABA-A and GABA-B are completely absent a steep linear function relates stimulus input and calcium signals, very similar to the way it happens in the calcium imaging experiments when PTX and CGP where added (figure 4B. cocktail). The modulatory effect was evident from very low stimulus intensities, which didn’t happen when one of both, GABA-A or GABA-B dependent conductances, was enabled. This phenomenon might emerge from the design of the network in which LNs do not inhibit only PNs but also other LNs. When GABA-A is blocked, LNs do receive less inhibition and thus, GABA-B mediated inhibition to PNs is enhanced. Similarly, when GABA-B is blocked, GABA-A mediated inhibition into PNs is enhanced. Such compensation might explain the observation that GABA-A or GABA-B components alone resulted enough to reduce the gain at the lower part of the function. Interestingly, model and experiments showed similar results in this regard, thus providing further indications that the AL model captures the key features of the real AL.

### Gain modulation and odor invariance

In the previous section we established that GABA modulates the gain function that relates input and output activity in the AL. However, to which extent does GABA contribute to stabilize the glomerular pattern that encodes the odor across concentrations has not been established before. Here we took advantage of the calcium imaging technique that allowed us to identify the same set of glomeruli across all bees and analyzed the stability of the glomerular pattern elicited by different concentrations of the same odor with and without the GABergic modulation. The analysis of pattern invariance was based on Pearsońs correlation coefficients between the glomerular activity patterns elicited by the different odorant concentrations. Correlations were calculated within each animal and between the activity pattern elicited by all concentrations excluding measurements with only-mineral oil. The comparisons among the 9 odor concentrations provided 36 correlation values within the concentration series measured with saline solution and 36 values within the concentrations series measured with the blockers. The table in figure 5A includes those 72 correlation values calculated in a representative bee. The upper half-matrix corresponds to the concentration series measured under control conditions and the lower half-matrix correspond to the concentration series measured in the same bee but under the effect of GABA-blockers PTX+CGP. The matrix is color-scaled to highlight the different correlation values. A glance at the colors distribution indicates that blocking GABA reduces the stability of the patterns, especially among patterns elicited by distant odorant concentrations. The scatterplot in figure 5B provides an overview of all correlation values along the first and the second series of measurements obtained for all bees. In this graph, the correlation obtained for a given pair of concentrations measured during the second series is expressed as function of the correlation determined for the same pair of concentrations during the first series. If the correlation values were similar, the point should not deviate from the expected line. As observed, a large proportion of pairs in the GABA-blockers group (purple dots) are below the line, consistent with lower correlations when GABA receptors were blocked. The table 5C, corresponds to the same matrix shown in 5A, but color-coded according to the relative distance between the two concentrations that are compared in each cell. For example, red cells indicate correlation values between pairs of patterns elicited by concentrations that are separated by a ∼3x factor from each other (i.e. dilution 1/3 vs. 1/10; 1/100 vs. 1/300; etc) while green cells indicate the correlation values between pairs of patterns elicited by concentrations separated by a 100x factor (i.e. dilution 1/3 vs. 1/300; 1/100 vs. 1/10000; etc). This distinction allowed us to pool correlation values based on the relative distance between the two concentrations that are being compared, and regardless of their absolute concentration. Figure 5D shows the average and SEM of correlation values for 9 bees treated with physiological saline solution during the first concentration series (black line) and with the PTX+CGP cocktail during the second series (purple line). GABA blockers reduced significantly the correlation coefficients between patterns elicited by odor concentrations that are separated by a factor of 30x or larger, thus supporting that GABA contributes to stabilize odor patterns.

**Figure 5.**
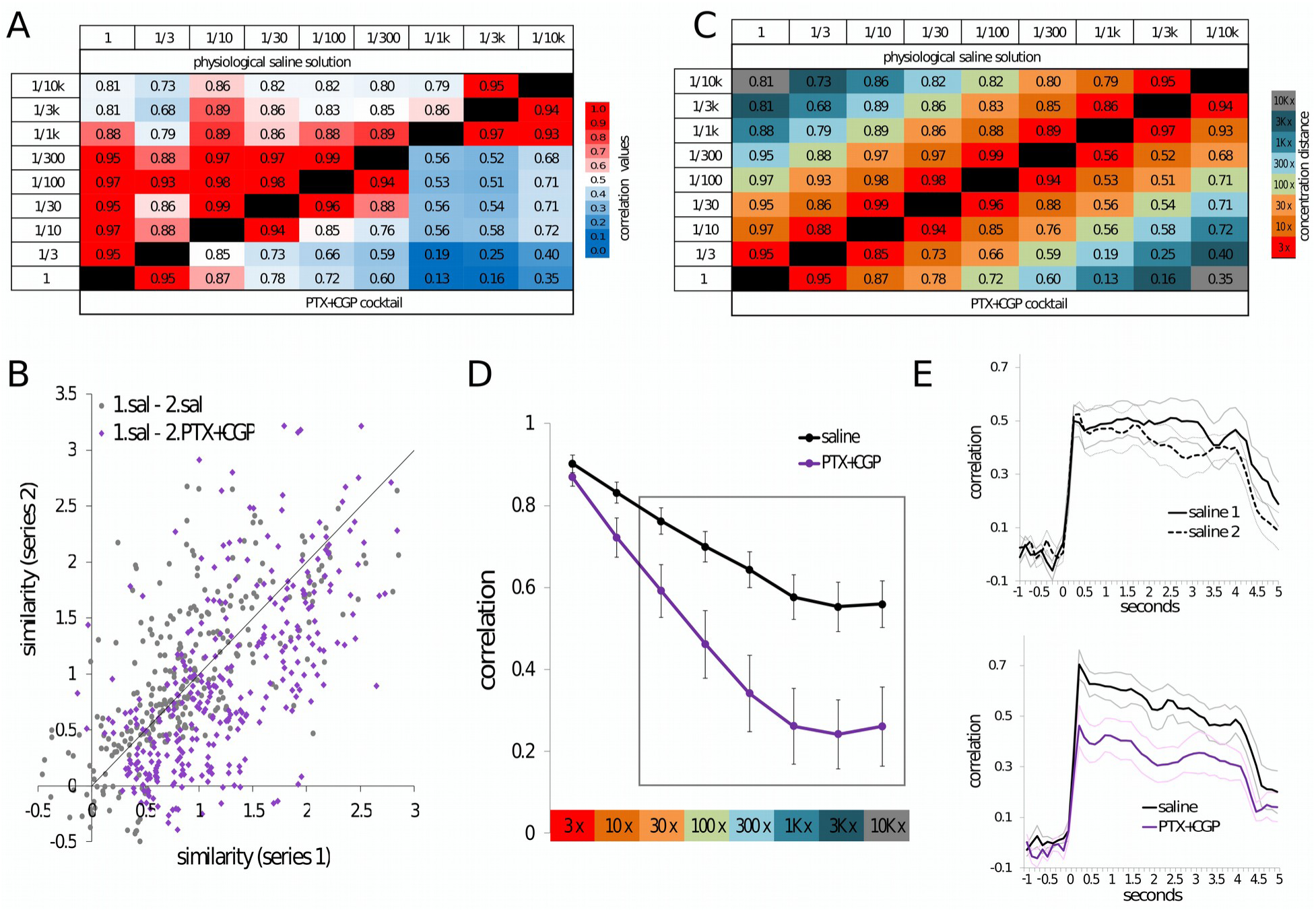
GABA and stability of the glomerular pattern. A. Cross-concentrations correlation matrix in a representative bee. Upper half matrix shows the correlations among glomerular patterns obtained during the saline series. Lower half matrix corresponds to the same bee but to the correlations among the patterns were obtained during the GABA blockers cocktail series. Color scale indicates the Pearsońs correlation value. B. Correlation values obtained for all possible pairs of patterns during the first and the second concentrations series for all bees measured in both series under physiological saline solution (gray dots) or measured first under saline and second under GABA blockers (purple). Correlation values are Fischer transformed and expressed as similarity index C. Same matrix as in A but colored to identify boxes that correspond to correlations among concentration pairs that keep the same relative distance. D. Mean and SEM of the correlations among concentrations pairs categorized as indicated by the matrix in C. Black: concentration series under physiological saline solution; purple: GABA blockers cocktail. The framed region indicates the points that are statistically different between both curves. E. Correlation values on a frame by frame basis among patterns within the first and within the second concentration series for bees treated with saline in both series N = 8 (upper panel) or treated first with saline and second with GABA blockers cocktail N = 9 (lower panel). Only odor pairs separated by a distance of 30x or larger were averaged, N = 11 bees.

The correlation coefficients shown from 5A to 5D, are based on the activity measured 375 ms after odor onset. Figure 5E extends the analysis to every 125ms acquisition frame, from one second before odor onset until one second after odor offset. All correlation coefficients between patterns elicited by odor concentrations that were separated by a factor of 30x or larger were averaged. As expected, signals are not correlated before odor onset, which is consistent with basal noise. 375 ms after odor onset, the correlation reaches its maximum value. The upper panel in figure 5E, shows the results obtained in bees that underwent the first and second series under physiological saline solution. The lower panel corresponds to bees that underwent the second series under GABA-blockers. As observed, the cross-concentrations correlation is affected during the whole stimulus duration, indicating that GABA-A and GABA-B dependent inhibition plays a critical role in stabilizing odor elicited activation patterns across intensities.

### Gain modulation increases with the number of active glomeruli

Previous studies in bees have shown that the number of active glomeruli increases with odor concentration (Sachse & Galizia, 2003). This may suggest that higher odor concentrations could lead to greater overlap in the glomerular patterns caused by different odors and therefore affect discrimination. However, it has been shown that honey bees discriminate odors better when odor concentration is high (Pelz et al., 1997; Wright & Smith, 2004). Consistent with this,Strauch et al., (Strauch et al., 2012) have shown that increasing odor concentration does not compromise pattern separation in the AL. These results indicate that gain modulation in the AL may not only favor pattern invariance across concentrations but also contribute to reduce the expected overlap when different odors increase their concentration. Here, we describe the relation between the number of recruited glomeruli and odor concentration, and test if this relation depends on GABAergic inhibition. For this analysis, we considered a given glomerulus as active when its average activity during one second after odor onset was two times higher than the SD of the activity measured before odor onset. Figure 6A shows that the number of recruited glomeruli increases with odor concentration and reaches a maximal value near to 60% of glomeruli. Instead, almost 100 % of glomeruli were activated when GABA receptors were blocked, which means that without GABAergic inhibition, no glomerular coding would be possible at high odor concentrations.

**Figure 6.**
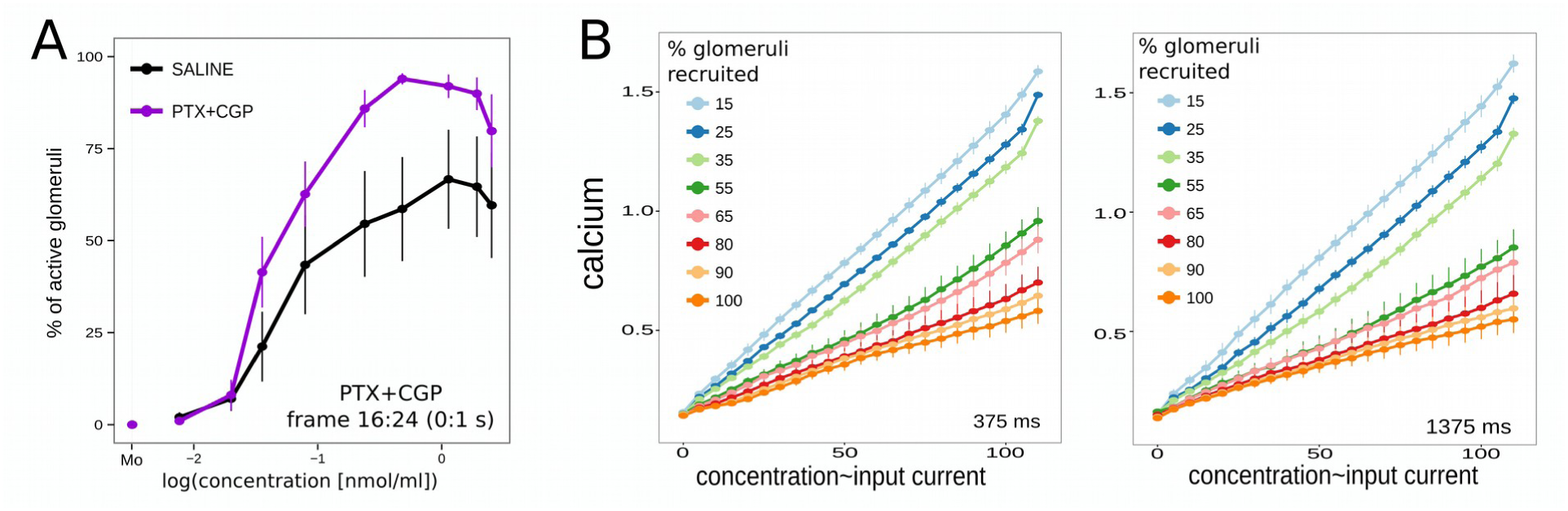
GABA and recruited glomeruli. A. Mean and SEM of the percentage of glomeruli recruited by different odor concentrations. Data corresponds to the group of bees treated first with physiological saline solution (black) and second with the GABA blockers cocktail (purple) N = 9. One glomerulus was considered as recruited when its activity during odor stimulations was two times larger than the standard deviation of the spontaneous activity during one second before odor onset. B. AL model output as function of injected current and number of recruited glomeruli. Data represent Mean and SEM of the intracellular calcium determined in active projection neurons, 375 and 1375 ms after odor onset.

Finally, the AL model gave us the opportunity to evaluate the interplay between the number of glomeruli that are recruited and the power of the GABAergic network that keeps odor elicited activity within a range that makes the glomerular code possible. In the previous sections (Figure 4c), different odor intensities were simulated by injecting different amounts of current to the same number of glomeruli. This time we tested the behavior of the AL network varying the amount of current and the number of glomeruli into which this current is injected. Figure 6B shows the average response of the PNs as function of the current injected and the percentage of recruited glomeruli. It is clear from the data, that when the proportion of recruited glomeruli increases, the slope of the gain function decreases. Interestingly, this is an experiment that can be made using only the AL model, since manipulating independently current and number of recruited glomeruli cannot be easy achieved in a biological system. In summary, the results indicate that the involvement and power of gain control is proportional to the number of recruited glomeruli.

## Discussion

Here we addressed the contribution that signal-processing in the antennal lobe does to keep odor representation relatively stable across concentrations. We describe the contribution that GABA-A and GABA-B receptors dependent-inhibition plays in terms of the amplitude and temporal profiles of odor elicited signals that convey information from the antennal lobes to the mushroom bodies. We found that GABA-A and GABA-B dependent inhibition increase the correlation among glomerular activity patterns elicited by different concentrations of the same odor. Using a realistic computational model of the antennal lobe we were able to run simulations that extend our characterization of the AL network by testing conditions that exceed experimental possibilities. Interestingly, even though based on rather simplistic connectivity rules and interactions among cells solely mediated by GABA-A and GABA-B interactions, the AL model was able to reproduce key features of the AL in relation to the stability of odor representation at different odor concentrations and decorrelation of projection projections neurons. Thus, the model provided a usable computational solutions that are of interest for the design of bio-mimetic artificial systems used in object recognition technologies.

### GABA: amplitude and temporal decorrelation of odor elicited activity

Fast and slow inhibitory interactions between antennal lobe neurons contribute to spike synchronization and slow temporal activity patterns that characterize odor responses at projection neuron level (Bazhenov et al., 2001; MacLeod & Laurent, 1996). In this framework, picrotoxin and CGP54626 have been used to selectively block GABA-A and GABA-B dependent inhibitory components in the antennal lobe of cockroaches and flies (Warren & Kloppenburg, 2014; R. I. Wilson & Laurent, 2005), as well as in the olfactory bulb of mice (McGann et al., 2005; Pírez & Wachowiak, 2008). Using these two blockers, here we show that GABA-A and GABA-B receptors shape odor responses in projection neurons of the honey bee AL. Measured by calcium imaging, GABA-A receptors produce a fast and transient inhibition of odor elicited signals that peaks 375ms after odor onset, and GABA-B receptors produce a slower and sustained inhibition that peaks 1375ms after odor onset. It has to be considered, that the temporal profile and magnitude of the signals measured in PNs correspond to the result of a complex interplay between excitatory input received from sensory neurons and inhibitory input received from local neurons, which in turn integrate excitation and inhibition from the same two sources (R. I. Wilson & Laurent, 2005). We designed a computational model of the antennal lobe to test the extent in which a neuronal network with this architecture and solely based on GABA-A and GABA-B dependent inhibitory interactions among its neurons is enough to generate spatio-temporal patterns of activity similar to those registered in the calcium imaging experiments. In the model, odor stimulation was simulated by injecting the same pulse of depolarizing current to all neurons in the recruited glomeruli. In spite of the uniform input pattern, several characteristic features of the AL output pattern, *i.e* spike synchronization, different intensities and diverse temporal profiles of the PNs responses were successfully obtained at the output side. Removing the currents associated to GABA-A receptors provided output patterns similar to those observed when the honey bee AL was treated with picrotoxin, *i.e.* an abrupt peak of activity shortly after odor onset and reduction in the correlation between glomeruli along odor stimulation (figure 3B). On the other hand, when currents associated to the GABA-B receptors were removed, the model provided output patterns similar to those produced by the honey bee AL under perfusion with CGP54626, *i.e.* different intensity responses across glomeruli that remained relatively stable during the whole odor stimulation (Figure 3D). When both, GABA-A and GABA-B dependent currents were enabled, the model mimicked realistic output patterns; *i.e.* the initial abrupt peak was attenuated, likely by the action of GABA-A receptors, and after that, the glomerular pattern evolved during odor stimulation with some glomeruli alternating their relative intensity, likely by the action GABA-B-dependent inhibition (figure 3E). This result supports what we know about the receptors associated with GABA-A (ionotropic) and GABA-B (metabotropic) and how the temporal dynamic of the two are different, GABA-A having a faster effect than the inhibition mediated by GABA-B. We are aware that other possible interactions already described among cells in the AL were no included in the present model, *i.e.* local neurons with different arborization patterns (Abel et al., 2001; Girardin et al., 2012), excitatory local interneurons (Shang, Claridge-Chang, Sjulson, Pypaert, & Miesenböck, 2007), electrical coupling among LNs and PNs (Huang, Zhang, Qiao, Hu, & Wang, 2010; Yaksi & Wilson, 2010) and inhibition pre-synaptic to the AL (Olsen & Wilson, 2008; Root et al., 2008). Interestingly, even though these interactions were not included, the simplified version of the antennal lobe was able to reproduce the main features of the AL responses. Remarkably, signal processing by the AL model, decorrelates the input channels, both in intensity and in time, two properties of the antennal lobe output that are linked to optimization of odor coding and separability of odor representation (Bhandawat, Olsen, Gouwens, Schlief, & Wilson, 2007; Silbering & Galizia, 2007). Interestingly, these properties were also shown in sensory processing and coding of other modalities such as vision (Hurvich & Jameson, 1957) and has been proposed as a convenient mechanism in terms of information theory (Buchsbaum & A Gottschalk, 1983).

### GABAergic modulation of the odor concentration-signal function

By doing calcium imaging of neural activity elicited by different odor concentrations under saline, PTX, CGP and cocktail conditions, we showed that GABAergic inhibition is crucial to attenuate the function that relates odor concentration and activity in PNs. The relationship between concentration and PNs activity was described by focusing on three main aspects of the signals: i) number of glomeruli recruited (fig 6.a); ii) intensity of calcium signals (fig 4.b), and iii) stability of the glomerular code (fig 5). In control saline conditions, the number of recruited glomeruli maintains a positive correlation with odor concentration until the three highest concentrations in which 60% of the glomeruli showed activity above detection threshold. When GABA receptors were blocked the relation was much steeper and almost 100% of the scrutinized glomeruli were active when stimulated with the five higher odor concentrations. The observation that 100% of the glomeruli showed activity when GABA receptors were blocked indicates that the value of 60% of glomeruli showing activity in saline condition, was not due to a limitation of the imaging technique neither a limit of responsiveness of the projection neurons, but rather due to a ceiling activity reached by the sensory input. Furthermore, it indicates that even saturating input levels can be handled by the GABAergic modulation to keep odor representation within the dynamic range of the PNs. The analysis of the intensity of the calcium signals in relation to increasing odor concentration was split to identify the contribution of GABA-A and GABA-B receptors. Interestingly, both receptors affect the gain function in different but partially overlapping concentrations ranges. Furthermore, the contribution of each one was evident during the time window determined as typical for GABA-A and GABA-B receptors. When a cocktail that blocks GABA-A and GABA-B receptors was used, the slope was affected along the whole concentrations range. Interestingly, we obtained in the AL model similar contributions of GABA-A and GABA-B to those obtained in the real AL. The gain function was affected by GABA-A receptors only during mid-to-high input levels, while the effect of GABA-B receptor was evident since very low input levels. When both receptors were functional a drastic modulation of the gain function was evident along the whole input range, suggesting synergistic functions of both mechanisms. Interestingly, the observation that GABA-A and GABA-B modulate the gain function at different concentration ranges emerges as a complex property of the network since no initial setting in the model dictates those differential sensitivities. Thus, these similarities validate the model and allow us to predict that similar emerging properties must determine the behavior of the real AL in relation to the contribution of the GABAergic inhibition to the gain modulation function.

### Gain function scaled to input intensity

As observed in the calcium imaging experiments, the higher the odor concentration, the higher the calcium signals amplitude and the number of glomeruli that were recruited. Because of this, in the last simulations, we studied the behavior of the model by co-varying both parameters (figure 6.b). It came out that the slope of the gain function scales down with the number of recruited glomeruli: the more glomeruli are recruited, the less the output signals are affected by the increase in the input. In this context, we can predict that low odor concentration recruits only few glomeruli and the inhibitory gain modulation plays no or a minimal role. However, higher odor concentration recruits more glomeruli and the inhibitory gain control is scaled accordingly to keep odor representation within a dynamical range. This phenomenon is consistent with previous studies in *Drosophila*, which demonstrated that the activity measured at single projection neurons scales in inverse proportion with the number of different receptors that are coactivated by the odor (Olsen et al., 2010).

Positive gain modulation in case of low odor concentrations has been shown in other models (Shang et al., 2007; Yaksi & Wilson, 2010; Zhu et al., 2013). A set of local neurons that form electrical and chemical synapses equalize concentration-invariant representations by spreading excitation or inhibition depending on the input strength. By solely modulating GABA we did not find evidences of positive gain modulations in case of low concentrations. However we found that GABA-A and GABA-B contribute to modulate the gain in partially different concentrations ranges. The existence of excitatory local neurons or electrical synpases in the honey bee AL still has to be investigated.

### Glomerular odor representations

It is well established that Kenyon cells, the post-synaptic partners of the PNs in the mushroom bodies, behave as decoders that detect specific patterns of co-activated projection neurons (Gruntman & Turner, 2013; Jortner, Farivar, & Laurent, 2007; Perez-Orive, 2004; Szyszka et al., 2005). In this context, it is expected that odor recognition is based on stable glomerular patterns, rather than on globally stable activity levels. The calcium imaging technique allowed us to identify the same set of glomeruli in all animals and focus the analysis on the stability of the glomerular pattern that encode the odor (figure 5). We found that blocking GABA receptors severely affected the correlation among glomerular patterns elicited by different concentrations of the same odor. This effect was specially marked among patterns of activity elicited by stimuli that differed in more than one order of magnitude in their concentrations (figure 5D). Previous studies in honey bees have shown that increasing odor concentration increases the separation among patterns of activity elicited by different odors, and that generalization between different concentrations of the same odors is possible thanks to smooth transitions among the patterns elicited by different concentrations (Sachse & Galizia, 2003; Stopfer et al., 2003; Strauch et al., 2012). Furthermore, evidences from the present and previous works indicate that local computations based on the GABAergic network accomplish these two properties. How is it that a mechanism based on GABA does reduce generalization in one case and increase generalization in another? Increasing odor concentration of an odor tends to broaden the pattern of receptors and glomeruli that are recruited. In case of two different odors, broadening their respective patterns might drive to partial overlap and a reduction in discrimination. Modulation of the gain function driven by GABAergic lateral inhibition keeps activation restricted to a core array of glomeruli for each odor minimizing overlaps and improving discrimination. In case of different concentrations of the same odor, the pattern elicited by the higher concentration includes the set glomeruli already activated by the lower concentration, plus secondary glomeruli that might change the perceptual quality of the odor (Johnson & Leon, 2000). Considering that the GABAergic modulation of the gain function is stronger in higher than in lower concentration, this modulation moves the pattern elicited by the higher concentrations closer to the pattern elicited by the lower concentration, eventually improving generalization among both concentrations. Our results are in line with previous reports in regards to concentration-invariant odor representations in the vertebrate olfactory bulb (Cleland, Johnson, Leon, & Linster, 2007; Wachowiak, Cohen, & Zochowski, 2017) and the insect antennal lobes (Asahina, Louis, Piccinotti, & Vosshall, 2009; Sachse & Galizia, 2003). In all cases it has been indicated that lateral interaction among co-activated glomeruli allows stable representations across concentrations (Olsen et al., 2010). Interestingly, a recent study was able to show the input-output transformation of odor representations in the mice olfactory bulb, providing direct evidence that OB output maps maintain relatively stable representation of odor identity across concentrations, even though the input maps can change markedly (Storace & Cohen, 2017). Our experimental and modeling results are consistent with such a capability and a critical role of GABA in this function of the AL.

### AL model architecture

To our knowledge, this is the first report of an AL computational model that incorporates glomeruli as sub-compartments. All PNs and LNs within each glomerulus receive the same excitatory input. LNs do not inhibit PNs that stem from the same glomerulus, but propagate inhibition to all other glomeruli depending on a given probability and independently of what the rest of the LNs do (Girardin et al., 2012). The consequence is a highly interconnected network in which the probability that a given glomerulus does not receive inhibition from another glomerulus is as low as 0.32 and the average number of LNs received from each glomerulus is 1 but could span between 0 and 5, thus providing different and random inhibitory strengths. LNs inhibit PNs but can also indirectly disinhibit them by inhibiting other LNs (Christensen, Waldrop, Harrow, & Hildebrand, 1993; Rein, Mustard, Strauch, Smith, & Galizia, 2013). In this way, the total amount of inhibition received by a PN in a given glomerulus is directly related with the number of glomeruli recruited by the odor and inversely related with the excitatory input received into this glomerulus, given that LNs that receive excitatory input in the same glomerulus can inhibit third part LNs. The lack of self-recurrent inhibitory feedback makes that only activation patterns that include two or more glomeruli are subject to inhibitory gain modulation. Activation patterns that include only one glomerulus are not affected by GABA, unless the odor concentration is so high that it starts recruiting secondary glomeruli. As it can be deduced from the architecture of the model network, inhibitory feedback scaled to input levels is based on the existence of odors that elicit multiglomerular patterns and exists in form of reciprocal inhibitory interactions among co-activated glomeruli, rather than on self-recurrent inhibitory feedback within each glomerulus. This design is in general terms consistent with the way in which non-pheromonal odors are detected and encoded, as co-activation of parallel channels (Hallem & Carlson, 2006), and later decoded by the Kenyon cells in the mushroom bodies (Gruntman & Turner, 2013; Jortner et al., 2007). Detection of pheromonal odors by specific receptors (Wanner et al., 2007) encoded by dedicated labeled lines (Sandoz, 2006) would be compatible with this network since they would actually benefit from having no recurrent inhibitory feedback. However, there are a number pheromones that are encoded as general odors and whose identity is recognized by the specific pattern of co-activated sensory channels (J. Carcaud, Giurfa, & Sandoz, 2015). Those patterns should not remain subordinated to the representation of general concurrent odors. In this context it is expected that the network established by the iLNs cannot be purely randomly distributed as it was set in the model, rather it should incorporate asymmetrical interactions that allow a hierarchical processing of the olfactory information.

### Partial and distributed gain control

Gain control at the antennal lobe or the olfactory bulb is not absolute (this work, and (Cleland et al., 2007; Olsen et al., 2010; Sachse & Galizia, 2003; Storace & Cohen, 2017; Strauch et al., 2012; Zhu et al., 2013). The transformations that take place between input and output patterns attenuate the effect of the concentration but do not make it completely independent. This was evident in our calcium imaging recordings (fig. 4). Furthermore, we could not find in the AL model a distribution or balance of inhibitions that provided absolute gain control. In this sense, our model and experimental results were consistent again.

In a way, it is reasonable that information on the concentration should not be completely lost, since different concentrations may be linked to different or even opposite values and eventually trigger different behaviors (Semmelhack & Wang, 2009). A solution to code quality and quantity without mixing them has been proposed by studies in honey bees and other Hymenoptera species. Honey bees, poses two parallels pathways of projection neurons that connect the ALs with the mushroom bodies (C. Giovanni Galizia & Rössler, 2010; Zwaka, Münch, Manz, Menzel, & Rybak, 2016). These pathways are the lateral antenno-protocerebral tract (l-APT), the same one that we have measured here, and the medial antenno-protocerebral tract (m-APT) (Kirschner et al., 2006). Interestingly, it has been described that PNs in the lAPT and mAPT may encode different properties of the olfactory information (J. Carcaud et al., 2015; C. Giovanni Galizia & Rössler, 2010; Hong & Wilson, 2015; Schmuker et al., 2011). In fact, Yamagata et al (Yamagata, 2009) found that lAPT PNs encode more efficiently odor identity while mAPT PNs are more sensitive to odor intensity. These determinations seem to contradict our description (also see (Sachse & Galizia, 2003)) of the effect of concentration on odor representation in lAPT PNs. The differences might be caused by the different recording sites. While Sachse et al (Sachse & Galizia, 2003) and we measured odor elicited activity in PNs dendrites in the AL, the determinations by Yamagata et al (Yamagata, 2009) were obtained from pre-synaptic boutons between PNs and Kenyon Cells at the mushroom body calyces. In this region, PNs boutons receive further GABAergic inhibitions as part of the local computation (Ganeshina & Menzel, 2001; Szyszka et al., 2005) as well as recurrent GABAergic feedback from the mushroom bodies output lobes (Grünewald, 1999; Rybak & Menzel, 1993). Thus, the partial gain modulation here reported in the AL must be interpreted as part of sequence of transformations of the olfactory information that continues along the olfactory pathways to the mushroom bodies (Froese et al., 2014).

### Gain modulation and plasticity

Notice that we show that the gain modulation function depends of the inhibitory synapses strength. Interestingly, we have shown in previous studies that activity dependent changes in the strength of the inhibitory synapses among AL neurons drive associative and non-associative plasticity linked to different types of learning (Chen et al., 2015; Locatelli et al., 2013). Now, based on the present results, we have to take into account the weight that those changes would have on gain modulation, likely improving intensity-invariant representation of learned odors. In this way, complementary functions of the AL network, such as gain-control and detection of relevant components might arise as emergent properties at the same time and by the same mechanisms for learned odors, providing animals the ability to detect and recognize meaningful odors presented at varying concentrations and in noisy backgrounds.

## Material and Method

### Experiments

#### Animals

Honey bee *Apis mellifera* pollen foragers were collected at the entrance of regular hives located at the campus of the University of Buenos Aires (34o 32’ S; 58o 6’ W). The bees were briefly cooled and restrained in individual holders suited for brain imaging recording (C Giovanni Galizia & Vetter, 2004). After recovery from cooling, bees were fed 5 *μ*L of a 1.0 M sucrose solution and remained undistributed until feeding *ad libitum* in the evening. While at the laboratory, bees were kept in a humid box at room temperature (20-24 °C) on a 12:12 h light:dark cycle. All experiments started 1 day after capture.

#### Odor Stimulation

The odor used was 2-octanone (TCI America, Portland OR). The odor delivery device, from now on the “odor-gun”, was conceived to provide 9 concentrations of 2-octanone or clean air upon demand. The odor-gun consisted of ten independent odor channels, each of them attached to 5 ml glass vial that contained the odorant diluted at different concentration. Each vial had a liquid volume of 300 µl. The dilutions in the liquid phase were in mineral oil: 1/10000, 3/10000, 1/1000, 3/1000, 1/100, 3/100, 1/10, 3/10 and pure octanone. A tenth vial contained only mineral oil. In all cases, the saturated headspace inside a vial was used as odor sample for stimulation. The odor-gun consisted in a central charcoal filtered air stream of 60 ml/s in which the headspace of one vial was pushed by nitrogen at 6 ml/s (nitrogen avoids odorant oxidation). Opening and closing of the odor channels were controlled by solenoid valves synchronized by the imaging acquisition software (TillVision). The final odor concentration delivered to the bee resulted 1/10 of the saturated headspace sample taken from the vials. During periods without odor stimulation the charcoal-filtered air stream ventilated continuously the antennae. An exhaust 10 cm behind the bee removed the odors.

#### Projection Neurons Staining

The head of the bee was fixed to the stage with soft dental wax (Kerr Sybron Dental Specialties, USA). A window was cut in the head capsule dorsal to the joints of the antennae and ventral to the medial ocellus. The glands were carefully moved aside until the mushroom body α-lobes were visible which serve as spatial reference for staining. Projection neurons (PNs) were stained by backfilling with the calcium sensor dye Fura-dextran (potassium salt, 10,000 MW; Invitrogen, Eugene, USA). The tip of a glass microelectrode coated with dye was inserted into both sides of the protocerebrum, dorsolateral of the α-lobes where the antenna-protocerebral tracts enter the lateral calyces of the mushroom bodies (Kirschner et al., 2006) (see Figure 1 first panel). The dye bolus dissolved into the tissue in 3-5 seconds. The window was closed using the piece of cuticle that had been previously removed and it was sealed with Eicosane (Sigma-Aldrich). Twenty minutes after staining, the bees were fed with 1 M sucrose solution and left undisturbed until next day. Before imaging, the antennae were fixed pointing towards the front using eicosane. The head capsule was opened and the brain was rinsed with Ringer solution (in mM: NaCl, 130; KCl, 6; MgCl_2_, 4; CaCl_2_, 5; sucrose, 160; glucose, 25; and HEPES, 10; pH6.7, 500 mOsmol; all chemicals from Sigma-Aldrich). Glands and trachea covering the ALs were removed. Only ALs that presented homogeneous staining of all visually accessible glomeruli were used for imaging. Body movements were prevented by gently compressing the abdomen and thorax with a piece of foam. A second hole in the head capsule was cut between the antennae and the mandibles, and the compact structure of muscles, esophagus and supporting chitin was lifted and put under slight tension. Finally, the brain was covered with Kwik-sil (WPI). After preparation, bees were mounted in the microscope and were allowed to recover for 20 minutes before starting with imaging.

#### Imaging

Calcium imaging was done using a EMCCD iXon camera (ANDOR, Belfast, UK) mounted on an upright fluorescence microscope (Olympus BX-50WI, Japan) equipped with a 20× dip objective, NA 0.95 (Olympus). Filter- and mirror-set: 505 DRLPXR dichroic mirror and 515 nm LP filter (Till-Photonics, Gräfelfing, Germany). Excitation light was provided by a Polychrome V (Till-Photonics) which alternated between 340 and 380 nm for excitation light. Acquisition protocols were made using the software TillVision (Till-Photonics). The sampling rate was 8 Hz and the spatial resolution was 125×125 pixels binned on a chip of 1000×1000 pixels. The intensity of the fluorescence lamp was controlled from the imaging acquisition software to get exposure times of 20 ms and 5 ms for 340 and 380 nm respectively. Each imaging session consisted of 20 measurements of odor-elicited activity (2 trials for each odor concentration). The odor concentrations were presented in increasing order and each trial was separated by 1 min intervals. Two series of increasing odor concentrations were performed. Both series were separated by 10 minutes giving time to change the perfusion solution. Each imaging measurement lasted 10 seconds and the odor presentation lasted 4 seconds, from second 2 to 6. The first concentration series was always performed under physiological saline solution. The second concentration series depended on the experiment: physiological saline solution, picrotoxin (Sigma), CGP54656 (Sigma) or cocktail of both blockers in all cases prepared in the same saline solution.

#### Pharmacology

The bees were divided in four groups, each treated with different solutions after and during the second odor concentration series: control bees (with saline), PTX (with picrotoxin GABA-A non-competitive antagonist) 10uM (Farkhooi et al., 2013), CGP54656 (GABA-B antagonist) 100uM (Warren & Kloppenburg, 2014; R. I. Wilson & Laurent, 2005) or a cocktail (PTX+CGP) 10uM + 100uM receptively.

#### Imaging Analysis

Imaging analysis was done using software written in IDL (Research Systems, CO, USA) by Giovanni Galizia (University Konstanz, Germany) and in R by Emiliano Marachlian. Each measurement consisted of two sequences of 80 fluorescence images each, obtained by alternating 340 and 380 nm excitation light (*Fi*_340_, *Fi*_380_, where *i* is the number of the image from 1 to 80). For each pair of image *Fi*, we calculated pixel-wise the ratio *Ri* = (*Fi*_340 nm_ / *Fi*_380 nm_) × 100 and subtracted the background *Rb*, obtained by averaging the *Ri* values 1 second before odor onset [*Rb* = 1/8 (*R*_8_ + … + *R*_16_)]. The resulting values (ΔR in figures) represent the change of fluorescence from the reference window and are proportional to the changes in the intracellular calcium concentration. The analysis of odor induced activation patterns in the present study was based on signals from 11 glomeruli that were identified on the basis of their morphology and position using published atlas of the honey bee AL (Flanagan & Mercer, 1989). Glomeruli are visible in the raw fluorescence images at 380 nm excitation light after backfilling the PNs with FURA (Figure 1B). In addition, we used a tool written in IDL by Mathias Ditzen (Freie Universitaet Berlin, Germany) that segments the image based on the degree of correlated activity between neighboring pixels. Pixels stemming from the same glomerulus are highly correlated. This provides images in which glomeruli are clear discrete units separated by dark boundaries (Figure 1B) and helps in the identification. The glomerular activation was calculated by averaging activity in a square area of 7×7 pixels that correspond to 23×23 *μ*m and fits within the size of the glomeruli.

### Modeling

#### Network Geometry

The antennal lobe model contained 20 glomeruli. Each glomerulus had 3 projection neurons (PNs) and 5 inhibitory local neurons (iLNs). All neurons (PNs and iLNs) in one glomerulus received the same excitatory input. The excitatory input and the number of glomeruli receiving it were varied across simulations to represent different stimulus intensities. The LNs constituted the network that interconnected glomeruli. Each LN received excitatory input only in one glomerulus but could inhibit every glomerulus with a probability of 25%. Thus, every LN inhibited an average of 5 glomeruli and every glomerulus received inhibition from an average of 25 LNs. LNs did not inhibit local PNs or itself, but could inhibit all other 4 LNs. The matrix in figure 2B shows a representative example of the connectivity between LNs to LNs and LNs to PNs in the network. Black squares denote inhibitory connections (the element in the column inhibits the element in the row).

The active glomeruli were selected based on a random function and we could change the probability of the glomeruli to receive excitatory input as a way to vary stimulus intensity. The excitatory input that simulated odor stimulation consisted in a depolarizing current that lasted 4 seconds with the form I=I_0_\**sc*\*exp(*rate*\*(t-t_0_)/1000), where I0 constitutes the parameter that was changed to vary the size of the sensory input, *sc* is the connection strength and *rate* corresponds to sensory adaptation (values in table 1). The input Io was varied across simulations between 0 and 100nA to represent different odor concentrations.

#### Model of Individual Neurons

Each PN and LN was modeled as two compartments (soma and axon) that included voltage and Ca^2+^ dependent currents based on the Hodgkin-Huxley model (Hodgkin & Huxley, 1952). Membrane potential of PNs and LNs was modeled by the two differential equations for the soma and axon respectively (1). Both contained leak and axon-soma coupled currents. The membrane potential equation for the soma included the calcium current *I_ca_*, the transient potassium current *I_A_*, synaptic GABA-A and GABA-B current *I_syn_*, a current that represented noise *I_r_* and the sensory input current *I*. The membrane potential equation for the axon included the sodium current *I_Na_*, the rectifier potassium current *I_kd_* and the potassium dependent calcium current *I_KCa_*.

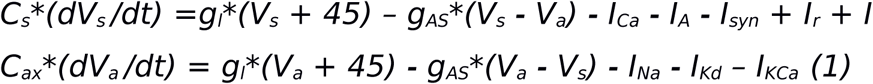

*C_s_* and *C_ax_* are the soma and axon capacity constants respectively (The values of all parameters are showed in table 1). *V_s_* and *V_a_* are the membrane potential of soma and axon membrane respectively. The two first terms to right of the equation are lineal. The conductances g_l_ and g_AS_ are constants (values in table 1). The first one corresponds to leak currents, and the second one is the axon-soma coupling term. The *I_ca_, I_A_, I_Na_, I_Kd_* and *I_KCa_* are described by the general equation (2).

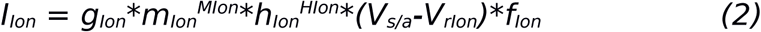

The subindex *Ion* indicates that this parameter or variable is ion specific. The gating variables *m* and *h* are temporal variables each one with its own dynamic equation (3). Each ionic current has its own gating variables and in the equation (2) are raised to different powers *M* and *H* (the values are in table 1). *V_s/a_* indicates the adequate soma or axon membrane potentials respectively *(V_s_* for *I_C_*_a_ and *I_A_* and *V_a_* for *I_Na_,I_Kd_* and *I_KCa_*) and *V_rIon_* is the rest potential for specific ion (table 1). *f_Ion_* is 1 in all currents except in *I_Ca_* in which *f_Ca_* is 1/(1+exp(2*Vs/24,42002442)).

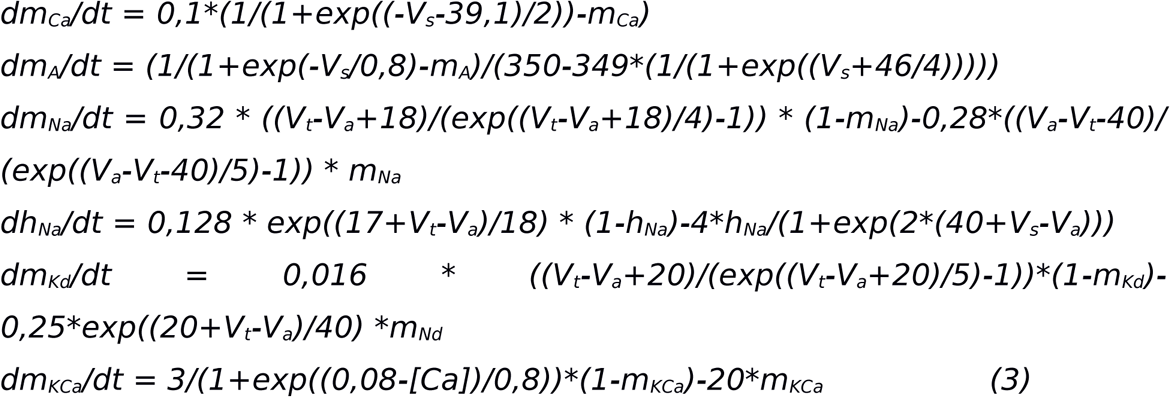

*V_t_* is a threshold membrane potential (show table 1) and [Ca] is the calcium concentration. For all cells, intracellular Ca^2+^ dynamics was described by a simple first-order model (4). *MU* is the calcium dynamic dissipation (values in table 1).

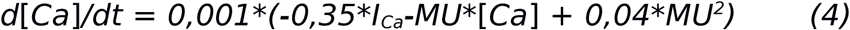

#### GABA Synaptic Currents

The only one connection between neurons considered in the present model was the GABA dependent synaptic current. We studied the action of both, GABA-A and GABA-B, dependent currents. Both are the result of LN activity but each one has its own dynamic. The synaptic current *I_syn_* was calculated as the summation of GABA currents *I_GABA-A_* and *I_GABA-B_*.

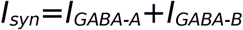

The GABA-A dependent current for the neuron *i* (*I_GABA-A_^i^*^)^ was given by the equation (5)

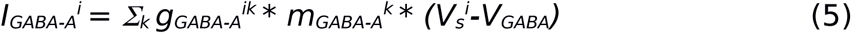

where *k* represents the LN that synapses with the *i* neuron and *g ^ik^* is the strength of the connection as in the figure 2B-C (for example, 0 if the *i* neuron was not connected with the *k* neuron). The gating variable m_GABA-A_ was varied in time and depended on its own dynamic equation (6) and *V_GABA_* is the resting membrane potential for GABA (value in table 1).

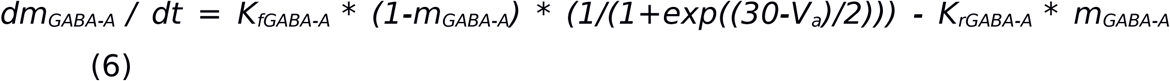

The values of *K_fGABA-A_* and *K_rGABA-A_* are in table 1.

The GABA-B current for the neuron *i* (*I_GABA-B_^i^*) was given by the equation (7)

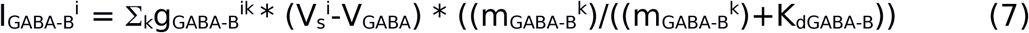

where *k* indicates the LN neuron that synapses with the *i* neuron. g_GABA-B_^ik^ is the value of strength connection setting as in the figure 2B-C. The gating variable m_GABA-B_ is temporal variable with its own dynamic equations (8) and *K_dGABA-B_* is a constant (value in table 1).

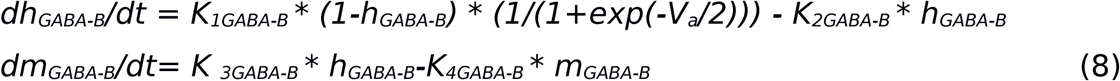

where *h*_*GABA-B*_ is an internal temporal variable.

The values of *K*_*1GABA-B*_, *K*_*2GABA-B*_, *K*_*3GABA-B*_ and K_4GABA-B_ are in table 1.

| Parameter | Symbol | LNI | PN |
| --- | --- | --- | --- |
| Soma capacity | $C_s$ | 10,0 nF | 10,0 nF |
| Axon capacity | $C_{ax}$ | 10,0 nF | 10,0 nF |
| Leak conductance | $g_l$ | 0,16 $\mu S$ | 0,16 $\mu S$ |
| Couple axon-soma conductance | $g_{AS}$ | 10,0 $\mu S$ | 65,0 $\mu S$ |
| $I_{Ca}$ powers values<br>Ca and rest potential | $M_{Ca}$ | 3 | 3 |
| | $H_{Ca}$ | 0 | 0 |
| | $V_{rCa}$ | 0 mV | 0 mV |
| $I_A$ powers values<br>A and rest potential | $M_A$ | 1 | 1 |
| | $H_A$ | 0 | 0 |
| | $V_A$ | -60 mV | -60 mV |
| $I_{Na}$ powers values<br>Na and rest potential | $M_{Na}$ | 2 | 2 |
| | $H_{Na}$ | 1 | 1 |
| | $V_{Na}$ | 50 mV | 50 mV |
| I<sub>Kd</sub> powers values<br>Kd and rest potential | $M_{Kd}$ | 1 | 1 |
| | $H_{Kd}$ | 0 | 0 |
| | $V_{Kd}$ | -60 mV | -60 mV |
| I<sub>KCa</sub> powers values<br>KCa and rest potential | $M_{KCa}$ | 1 | 1 |
| | $H_{KCa}$ | 0 | 0 |
| | $V_{KCa}$ | -60 mV | -60 mV |
| Threshold potential | $V_t$ | -51,7 mV | -52,1 mV |
| Calcium dynamic dissipation | MU | 1,5 | 1,6 |
| Strength connecting | $Sc$ | 0,5 | 0,7 |
| Sensory adaptation | $rate$ | 0,05 | 0,05 |
| GABA rest potential | $V_{GABA}$ | -90 mV | -90 mV |
| | $K_{fGABA-A}$ | 10,0 ms<sup>-1</sup> | |
| | $K_{rGABA-A}$ | 0,1 ms<sup>-1</sup> | |
| | $K_{dGABA-B}$ | 100 | |
| | $K_{1GABA-B}$ | 0,6 mM<sup>-1</sup>ms<sup>-1</sup> | |
| | $K_{2GABA-B}$ | 0,001 | |
| | $K_{3GABA-B}$ | 0,3 | |
| | $K_{4GABA-B}$ | 0,0025 | |

